# PABPC1 Modulates Immunoglobulin pre-mRNA Alternative Polyadenylation

**DOI:** 10.64898/2026.04.23.720383

**Authors:** Zachary Miller, Jake Dearborn, Ramiro Barrantes-Reynolds, Hana Paculova, Drew Honson, Jihui Sha, Weixian Deng, Tylar Kirch, William Dowell, Sylvester Languon, Kalev Freeman, Seth Frietze, James Wohlschlegel, Devdoot Majumdar

**Author notes:** <u>Corresponding Author:</u> Devdoot Majumdar, <u>Email:</u>. <u>Author Contributions:</u> Author contributions were provided by CRediT taxonomy at the time of submission.

## Abstract

Alternative polyadenylation is a mechanism by which cells tune gene expression, and dysregulation can lead to development of disease. PABPC1 has been implicated in poly(A) site selection, but its function in gene regulation remains contradictory and poorly defined. Here, we investigate its role in B cell development, where APA controls immunoglobulin secretion. To define this role, we mapped PABPC1-RNA interactions using CLAP-seq and perturbed PABPC1 expression using a degron based strategy. PABPC1 localizes to the 3’UTR in 70% of its gene targets and primarily binds to A-rich regions. Integration with transcriptomic data suggests PABPC1 downregulates 60% of its gene targets. While transcriptome-wide shifts in 3’ UTR length were limited, PABPC1 binding was specifically enriched in genes exhibiting significant 3’ UTR shortening. Using a foundational genomics model, we find the PAS-proximal region is the most predictive of gene expression within PABPC1 binding sites. Positional analysis revealed PABPC1 localizes closer to the PAS in genes downregulated following depletion. In immunoglobulin transcripts, PABPC1 binds to both secreted and membrane isoforms and is more enriched at the secretory PAS, and depletion modestly alters immunoglobulin expression. Together, our findings demonstrate PABPC1 primarily shortens and downregulates its targets in a context dependent manner.

**Graphical Abstract:** 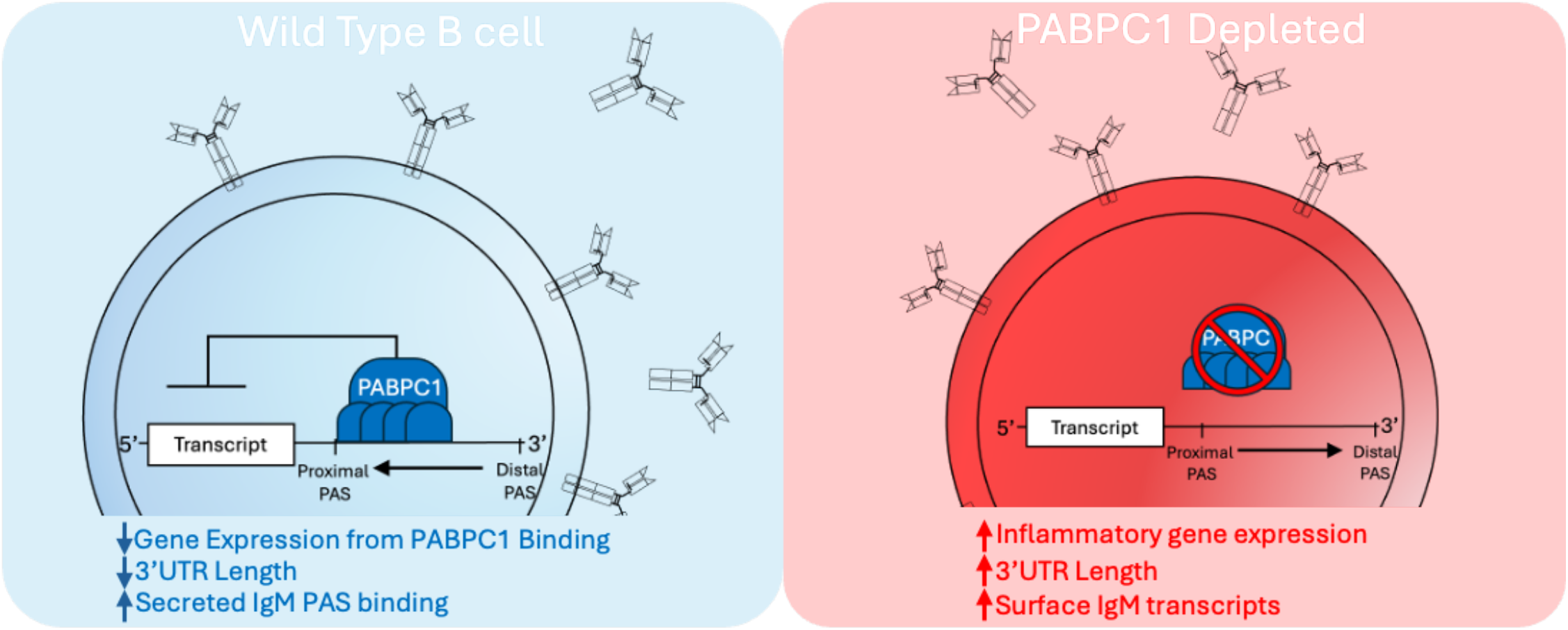

## Introduction

B cells sit at the center of the humoral immune response by producing highly specific antibodies, and their proper development underpins functional immune memory, immune tolerance, and pathogen clearance. These processes are conducted by virtue of precisely timed gene expression, where aberrant expression of crucial immune genes sometimes results in incomplete pathogen clearance, autoimmunity, and cancer (1–3). Importantly, gene expression is regulated by the polyadenylation machinery, which recognizes the conserved hexameric signal sequence (‘AAUAAA’) (4) in order to catalyze the subsequent cleavage and polyadenylation of a maturing pre-mRNA. As many as 70% of coding genes contain multiple poly (A) signal sequences (PAS) (5) and tightly regulated PAS selection can tune the length of mRNA through alternative polyadenylation (APA), which can result in drastic functional consequences on expression (6).

In the immune system, APA determines if an antibody (also known as immunoglobulin, Ig) is secreted (sIg) or expressed at the cell membrane (mIg) (7–9). Expression of mIg enables affinity maturation, whereas sIg expression enables an effective antibody-mediated immune response against antigens. The choice between these mIg and sIg is made by APA: the transmembrane domain that is necessary for surface expression is flanked by two PAS. In B cells, pre-mRNA cleavage and polyadenylation occurs at the distal PAS leading to mIg expression. However, when B cells terminally differentiate to plasma cells (PCs), processing at the proximal PAS leads to sIg expression. Therefore, precise regulation of APA in B cells is crucial to mounting an effective immune response.

While it is commonly understood that APA regulation determines whether an antibody is secreted, the mechanism by which PAS selection occurs in B cells is not completely understood. Early models for PAS selection suggested that CSTF64, a core CPA component, is limiting in B cells but not in PCs (10, 11). Modern transcriptomic studies, however, do not support dynamic regulation of CSTF64 throughout B cell development, although several studies suggest many indirect mechanisms (12–15).

One compelling model is that Cytoplasmic Poly(A) Binding Protein 1 (PABPC1) is able to traffic hnRNPLL to stimulate processing at the sIg PAS (15). In other contexts, PABPC1 interacts with the poly(A) tail to regulate transcript stability (16, 17), traffic fully processed transcripts between cellular compartments (18, 19), facilitate translation (20–23), and mounting evidence indicates it contribute to PAS selection in other systems (24–26). However, it remains unclear if PABPC1 interacts with Ig transcripts specifically or if the localization is mediated by its affinity to the poly(A) tail. The pleiotropic functions of PABPC1 defined in other systems coupled with conflicting reports on the precise role PABPC1 plays in regulating APA and gene expression (26–29) leave its function in B cells muddled and poorly defined.

Here, we investigate the role of PABPC1 in B cell development and function by integrating RNA binding sites and functional perturbation. Transcriptome-wide mapping revealed that PABPC1 predominantly localizes to the 3’ Untranslated Region (UTR). Following depletion, gene expression was primarily increased, indicating PABPC1 broadly represses transcript abundance. Loss of PABPC1 additionally resulted in lengthening of its target transcripts, consistent with a role in promoting use of proximal polyadenylation sites. To determine if these changes could be explained by cis-regulatory features, we correlated PABPC1 binding with expression changes and applied a foundational genomics model to these regions to identify sequences predictive of gene expression. PABPC1 binding sites within upregulated genes were more likely to contain the canonical PAS, whereas PABPC1 was positioned closer to the PAS in downregulated genes. Extending these observations to immunoglobulin transcripts, we found depletion of PABPC1 increased both mIg and sIg, while reducing total protein levels, highlighting its disparate roles in transcriptional regulation and translational control. Taken together, we find PABPC1 binding represses gene expression and promotes use of proximal poly-A sites. In B cells, our finding that PABPC1 promotes use of the secretory poly-A site may reflect that this regulator exerts its influence in a previously unrecognized, context specific way.

## Materials and methods

### Cell Culture

All experiment were performed using Ramos B cells. Cells were cultured at 37°C under 5% CO_2_ in RPMI-1640 media supplemented with 1x MEM non-essential amino acids, 1x Sodium Pyruvate, HEPES, 1x penicillin-streptomycin, beta-mercaptoethanol, and Bovine Calf Serum. CLAP experiments were performed in cells stably expressing doxycycline inducible HaloTag-RBP fusions. Degron experiments were performed using cells in which FKBP12F36V was knocked into the c-terminus of PABPC1.

### Generation of Cell lines

Genome editing and transgene integration were performed using the Neon electroporation system (ThermoFisher Scientific, Waltham, MA) with the following conditions: 1350V for 1 pulse with a pulse width of 30 ms. HaloTag fusion constructs were introduced using the piggyBac expression system. Coding sequences for PABPC1 and PTBP1 were amplified from Ramos cDNA and cloned using Gateway Recombination technology (ThermoFisher Scientific, Waltham, MA).

To establish degron cell lines, sgRNAs (Supplementary Table 1) targeting the PABPC1 C-terminus were cloned into lentiGuide-Puro. A homology-directed repair (HDR) template encoding FKBP12F36V-P2A-mCherry with roughly 500 bp homology arms flanking the PABPC1 stop codon was assembled using a Golden Gate cloning strategy into a domesticated pUC19 plasmid (Supplementary Table 1). The PAM sequences in the homology arms were mutated to prevent Cas9 re-cleaving the transgene. Correct integration was validated using western blot.

Lenti-Cas9-2A-Blast was a gift from Jason Moffat (Addgene plasmid # 73310; http://n2t.net/addgene:73310; RRID: Addgene_73310). pB-Halo,IRES-eGFP backbone, which was a gift from Mitchell Guttman (Addgene plasmid # 164519; http://n2t.net/addgene:164519; RRID: Addgene_164519). LentiGuide-Puro, which was a gift from Feng Zhang (Addgene plasmid # 52963; http://n2t.net/addgene:52963; RRID: Addgene_52963),

### Cross-Linking and Affinity Purification

Cross linking and affinity purification was performed as previously described by Guo et al. (34): Briefly HaloTag fusion proteins were induced with 2 μg/mL doxycycline for 16 hours. Cells were crosslinked with UV-C light on ice in a Stratalinker UV1800 (Agilent Tecnologies, Inc, Santa Clara, CA) in duplicate. Crosslinked cells were lysed and incubated with HaloLink Resin beads (Promega, Madison, WI) to pulldown RBP-bound RNA.

Beads were washed sequentially under stringent conditions at 91°C using buffers containing SDS, high salt concentration, urea, or NP-40. Protein was digested from RNA-Protein complexes with proteinase K, and RNA was eluted using Pierce microspin cups (ThermoFisher Scientific, Waltham, MA).

### Library Prep

RNA was purified at every step using Zymo Clean and Concentrate 5 kit (Zymo Research Operations, Tustin, CA). Input RNA was treated with DNase I (NEB, Ipswich, MA) and fragmented briefly at 91°C. RNA ends were repaired with FastAP (ThermoFisher Scientific, Waltham, MA) and PNK (Bayou Biolabs, Metairie, LA). Following purification, RNA was ligated to an adapter (Supplementary Table 1) containing a priming site for RT using high concentration T4 RNA ligase (NEB, Ipswich, MA). To exclude excess adapter, RNA was purified using the zymo >200nt protocol Reverse Transcription was performed using Protoscript II RT (NEB, Ipswich, MA). To degrade the RNA, samples were treated with NaOH at 80°C, and the second adapter (Supplementary Table 1) was ligated using T4 DNA ligase (Bayou Biolabs, Metairie, LA). Libraries were amplified using adapters specific to the Singular sequencing platform. Libraries were gel purified from fragments between 180 and 700 bp, pooled for sequencing, and sequenced using a Singular G4 sequencer at the University of Vermont Integrative Genomics Resource Core Facility.

### Bulk RNA-seq

Total RNA was extracted from 1E6 cells using the Zymo Quick-RNA Miniprep kit (Zymo Research Operations, Tustin, CA) according to manufacturer instructions. cDNA libraries were generated using template-switching RT primers using Protoscript II RT (NEB, Ipswich, MA) and cDNA was amplified and prepared for sequencing using the Nextera XT (illumina, San Diego, CA) kit per manufacturer instructions. Libraries were sequenced using the HiSeq Illumina sequencing platform.

Reads were aligned to a human reference genome (hg38) using STAR. Counts for each gene were quantified with FeatureCounts, and differential expression was determined using the DESeq2 package in R, using 0.585 and −0.585 as the log2FC cutoff and a significance cutoff of adjusted p value <0.05. Read Density Histograms were produced using Deeptools normalized by BPM with a bin size of 10bp and plotted using custom scripts. Labeled genes in the volcano plot are those that meet the log2FC and adjusted p value cutoffs and are part of gene ontologies that are related to the immune system. Immune genes were determined using MSigDB hallmark gene sets and Kegg pathways.

### Bioinformatic Analyses

CLAP-seq libraries were analyzed using the Skipper workflow, as described previously(35). Metagene profiles were generated using Homer2. Analysis of changing 3’UTR length was performed using DaPars2 and alternative splicing was quantified using rMATS-turbo. Enriched motifs were displayed graphically using ggseqlogo. Correlational analyses and data visualization was performed using custom R scripts. Gene set enrichment analysis was performed using the fgsea R package following DESeq2 quantification.

### Machine Learning

To detect patterns in close proximity to PABPC1 binding sites, we first selected genes with differential expression identified by DESEQ2 following PABPC1 depletion. We filtered those genes to include only regions with PABPC1 enriched binding windows, as identified in our CLAP-seq dataset. Next, we used Borzoi (91), a foundational genomics model that predicts gene expression from DNA sequences, to predict gene expression in naïve B cells. Features, or genomic regions, that the model used to predict expression, centered on PABPC1 binding windows within differentially expressed genes, were scored using saliency. TF-MoDISco (92) was then used to determine *De Novo* motifs based on the saliency score as regions predictive of gene expression (Fig 3G). To identify regions of interest, we included motifs that were within 50bp of the binding windows identified using CLAP-seq, which were generally 100bp wide. Motifs enriched in PABPC1 binding sites of upregulated genes were then compared to downregulated regions to determine relative enrichment using FIMO (93, 94).

### Western Blot

Cells were lysed in RIPA buffer and protein was quantified using Pierce BCA assay (ThermoFisher Scientific, Waltham, MA). Proteins (30-40ug) were resolved on Bis-TRIS polyacrylamide gels (GenScript Biotech, Nanjing, China). Following electrophoresis, protein was transferred to nitrocellulose. Membranes were blocked for 1 hr at room temperature in 5% Milk/TBST with agitation and incubated overnight with primary antibody against PABPC1 (Cell Signaling Technologies Antibody #4992) or GAPDH (Cell Signaling Technologies Antibody #2118). HRP-conjugated secondary antibody (Cell Signaling Technologies Antibody #7074) was incubated for an hour in 5% Milk/TBST with gentle agitation. Signal was detected using Immobilin Forte Western HRP substrate (Sigma-Aldrich, Merck Group, St. Louis, MO) per manufacturer instruction and chemiluminescence was detected using an Amersham Imager 600 (E, Boston, MA).

For validation of PABPC1 degradation following dTag13 treatment, cells were treated with either dTag13 (500 nM) or DMSO as a vehicle control for 16 hours in duplicate.

### RT-qPCR

RT-qPCR experiments, Luna Universal Master Mix (NEB, Ipswich, MA) was used, and performed on a QuantStudio 3 Real-Time PCR Instrument (Applied Biosystems, ThermoFisher Scientific, Waltham, MA). cDNA was generated using oligo-d (T) to prime the reaction, and Protoscript II RT (NEB, Ipswich, MA). Standard curves were established using gel-purified amplicons corresponding to sIg and mIg isoforms to enable absolute quantification. Primers used for RT-qPCR are detailed in Supplementary Table 1

For isoform analysis, cells expressing PABPC1-FKBP12F36V and wild-type controls were treated with 500 nM dTag-13 or DMSO for 16 hours. DNA copy number was calculated from the standard curves.

### Intracellular Flow Cytometry

Cells Expressing PABPC1-FKBP12F36V and wild-type Ramos were treated with 500 nM dTag13 or DMSO vehicle control for 16 hours. mIg was stained using Alexa647 conjugated anti-human IgM (Biolegend cat. #314536, San Diego, CA). Cells were fixed with 4% PFA, permeabilized using saponin and intracellular Ig was stained using FITC conjugated anti-human IgM (Biolegend cat. #314506, San Diego, CA). Samples were analyzed on a MACSquant 10 (Miltenyi, Westphalia, Germany) and processed using FlowJo (Tree Star, Ashland OR).

### Plasmid Validation

All plasmids were verified by whole plasmid sequencing from Plasmidsaurus using Oxford Nanopore Technology, or by Sanger Sequencing from Eurofins Genomics.

### Statistical Analyses

Data are presented as mean +/-standard deviation (S.D.) of at least n=2 biological replicates, as indicated in figure legends. Statistical significance for pairwise comparisons was determined by an unpaired, two-tailed Student’s t-test. For global miRNA distribution, a Wilcoxon rank-sum test was performed. Gene Set Enrichment Analysis (GSEA) was performed using rank gene lists. P < 0.05 were considered statistically significant.

## Results

### PABPC1 binds primarily in the 3’UTR of mRNA targets in Human B cells

Determining precisely where an RNA binding protein (RBP) binds can provide valuable insight into its function and mechanistic specificity. Previous studies have mapped PABPC1 using an immunoprecipitation based strategy, which established that it primarily binds A-rich regions within the 3’ Untranslated Region (3’UTR) (30– 32). Because PABPC1 has been shown to influence IgM secretion, but has never been directly mapped in B cells, we sought to first identify where it binds globally to understand the mechanistic role it plays in RNA processing in the adaptive immune system. To that end, we utilized cross linking and affinity purification coupled with sequencing (CLAP-seq). Where previous approaches have used antibodies to enrich RBP-RNA interactions, CLAP-seq involves fusing an RBP to a HaloTag. The covalent interaction formed between Halo and its ligand allows the use of high temperatures and concentrated detergents during purification to reduce non-specific interactions that are not possible with typical antibody-based strategies (33). PABPC1 was therefore fused to HaloTag and stably expressed in Ramos B cells before crosslinking and enrichment (Fig 1A). To account for the possibility of PABPC1 overexpression artifacts, we used a doxycycline inducible system, as previously described (34). We used the Skipper workflow (35) to identify PABPC1 binding windows significantly enriched over background. Importantly, concordance of binding regions between replicates was significant, indicating CLAP-seq can capture reproducible RBP binding regions (Fig 1B).

**Figure 1.**
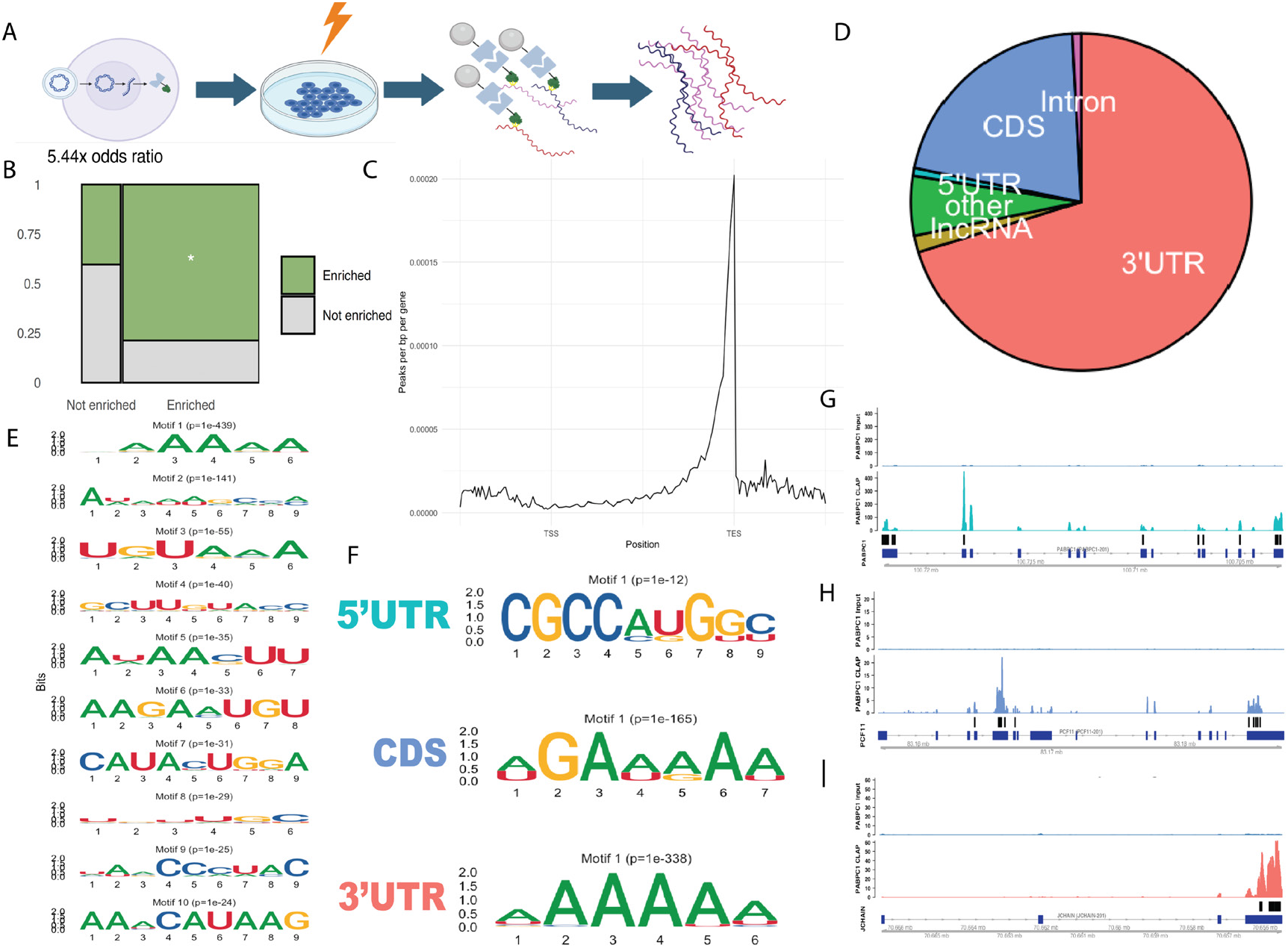
PABPC1 binds primarily A-rich motifs in the 3’UTR of B cells. **A** Schematic for CLAP-seq. Halo-tagged fusion proteins crosslinked to RNA are enriched using Halo sepharose beads, and enriched RNA is sequenced and compared input control to identify enriched regions **B** Concordance between enrichment of PABPC1 binding sites in replicates following CLAP-seq (N=2). Regions enriched in one replicate were 5.4 times more likely to be enriched in the other replicate, compared regions that were not enriched. **C** Metagene analysis of PABPC1 peak density over normalized gene length. Transcription start sites (TSS) and Transcription End Sites (TES) are marked as indicated. **D** Reproducibly enriched PABPC1 binding sites were sub-categorized to determine where in gene bodies PABPC1 binds in its targets. 70% of PABPC1 targets are in the 3’UTR, and roughly 20% fall in coding sequences. **E** Top ten motifs in PABPC1 enriched binding windows over background regions. The canonical poly(A) motif is the most significantly enriched (p=1E-439). **F** PABPC1 binding windows were categorized based on where they bind in targets. The most enriched motifs in the CDS and 3’UTR were A-rich, as expected, but in the 5’UTR, the most enriched motif was CGCCAUGGC, a motif that contains the mammalian Kozak sequence. Intronic motifs were excluded because the top motif did not meet the significance threshold to exclude false positives. **G** Representative read density histogram of PABPC1 enriched windows in the 5’UTR of PABPC1. **H** Representative read density histogram of PABPC1 enriched windows primarily in the CDS of PCF11. **I** Representative read density histogram of PABPC1 enriched windows in the 3’UTR of JCHAIN. **G-I** black bars indicate enriched regions as identified by Skipper.

PABPC1 was shown to bind A-rich regions primarily located in the 3’UTR in other cellular contexts (32). To determine if known PABPC1 binding patterns hold true for B cells, we performed metagene analysis of enriched binding regions using Homer (36). As expected, PABPC1 localized primarily to the 3’ when interacting with RNA (Fig 1C). When we subset binding windows based on location in gene bodies, we found 70% of significantly enriched PABPC1 binding windows reside in the 3’UTR, while 20% are in the coding sequences (CDS) of target mRNA (Fig 1D), consistent with the known binding patterns of PABPC1 (37). Analysis of the motifs within significantly enriched binding sites revealed that a large fraction of binding can be attributed to the sequence ‘AAAAA’ (p=1E-439, Fig 1E), consistent with the well-characterized interaction between PABPC1 and the poly(A) tail (38–42). Motif identification within binding sites subset by gene regions showed that in the 3’UTR and CDS, PABPC1 primarily bound to A-rich regions (Fig 1F), consistent with our global analysis (Fig 1E). Unexpectedly, the top motif in 5’UTR targets contained the mammalian Kozak sequence (43). This is likely because PABPC1 forms a closed loop structure with the RNA cap binding complex to facilitate ribosome scanning and translation. This may reflect a role for PABPC1 in stabilizing this interaction through interaction with the kozak sequence.

Like many RBPs, PABPC1 is known to autoregulate its expression by binding in the 5’UTR of its own mRNA (32, 44, 45). We found PABPC1 binds to the 5’UTR of its own transcript, as shown in other systems (Fig 1G). Because this regulation is mediated by direct PABPC1 binding, we reasoned PABPC1 binding in other genes may hint at regulatory role for those genes as well. Representative read density histograms of PABPC1 binding windows in the CDS and 3’UTR show PABPC1 binds directly to PCF11, an important APA regulator (46) and JCHAIN, a protein responsible for linking monomeric IgM and IgA in PCs (47) (Fig 1H-I). This suggests PABPC1 may influence expression of these genes in the B cell lineage. Together, our data represent the first high-resolution map of PABPC1-RNA interactions in B cells, revealing both conserved binding patterns and previously undefined roles to help immune gene regulation.

### PABPC1 modulates immune gene expression in B cells

B cell development and function result in proper elicitation of humoral immunity, immune memory, and proper discrimination of self-versus non-self by exacting precise modulation of timing and valence of hundreds of immune genes. Regulation of polyadenylation and recognition of PAS sits at the center of post-transcriptional regulatory pathways that help modulate immune gene expression, including mRNA production, trafficking, degradation, and protein production. After mapping PABPC1 in B cells primarily to the 3’UTR, we next considered that PABPC1 may influence gene expression. To understand the gene expression programs elicited by PABPC1 interaction, we sought to perturb its expression. Because PABPC1 is an essential gene (27, 48, 49) that is deleterious to cell health upon full knockout, we employed a degron based strategy that preserves endogenous gene expression while enabling rapid small molecule-mediated shut-off. Briefly, we utilized the dTag system by harnessing Cas9 to generate a C-terminal fusion of PABPC1 containing a FKBP12F36V, which facilitates dTag13 binding, as described previously (50). Upon dTag13 binding, the fusion protein is ubiquitinated and targeted for degradation (FIG 2A). We were able to generate several biallelic clones (Fig 2B), allowing acute control of PABPC1 and complete shut-off by 16 hours (Fig 2C). Of note, in knock-in lines we observe a faint band around 70 kDa, roughly equivalent in mass to endogenous PABPC1. This is likely PABPC3, which is expressed at lower levels than PABPC1, but bears 92% homology (51) and is recognized by our PABPC1 detection antibody. Importantly, to our knowledge, interrogation into the role of PABPC1 using functional genomics have been limited to siRNA (15, 26) or CRISPRi (28) modalities, and this represents the first acute perturbation method applied to PABPC1, evading challenges of cell line compensation and allowing for rapid depletion.

**Figure 2.**
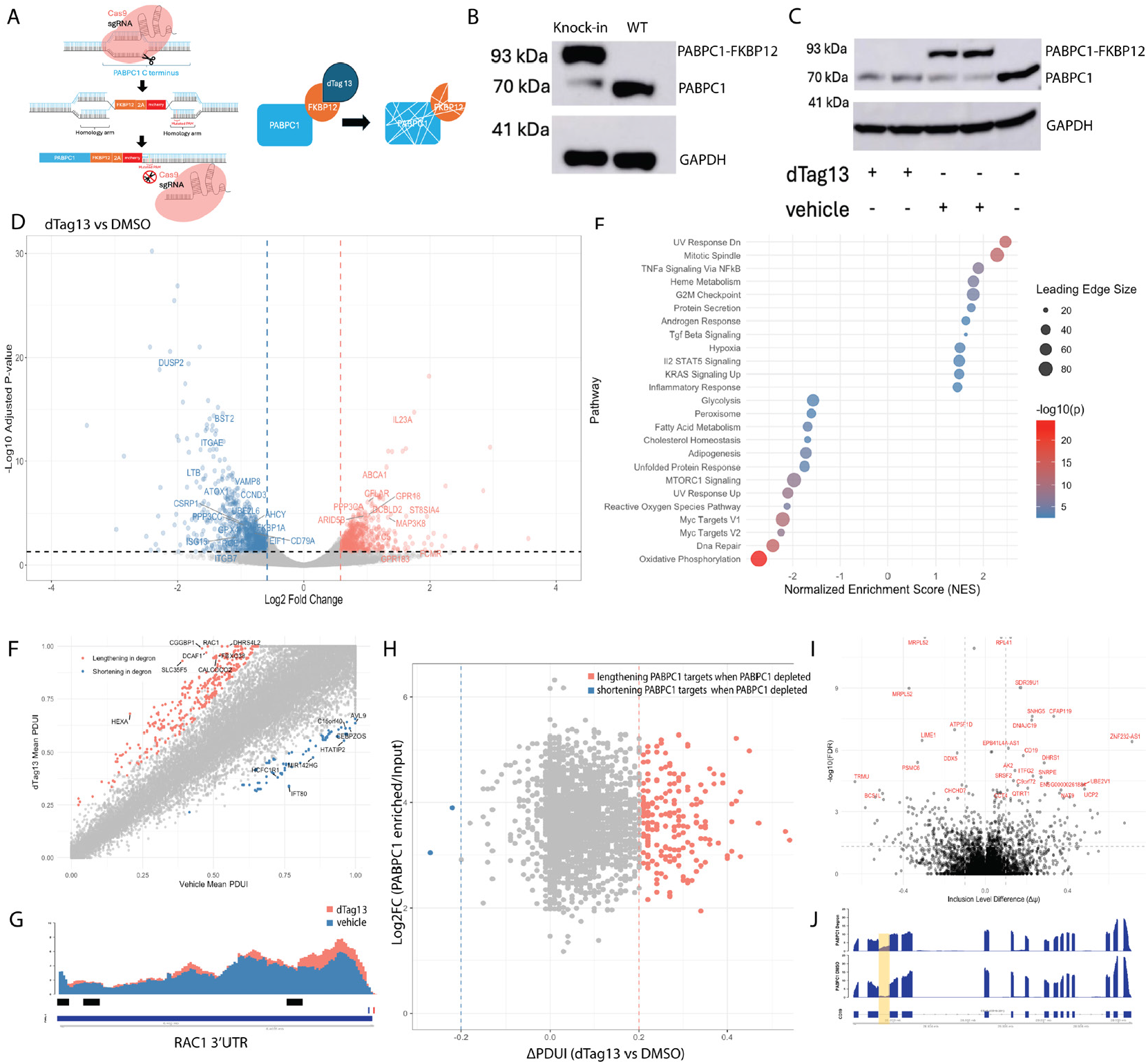
PABPC1 depletion alters immune gene expression, but only modestly changes 3’UTR length and alternative splicing in B cells. **A** Schematic depicting strategy to knock in the FKBP12 to the PABPC1 locus. **B** SDS-PAGE gel for PABPC1 following electroporation of guide RNA plasmids, Cas9, and HDR template. WT PABPC1 is expected to migrate to 70 kDa, and the fusion is expected to migrate to 93 kDa. GAPDH was used as a loading control. **C** SDS-PAGE gel of Ramos B cells expressing PABPC1-FKBP12 fusions treated with either dTag13 or DMSO for 16 hours. **D** RNA-sequencing was done on PABPC1 depleted cells following 16 hours of dTag13 treatment (N=2). Genes upregulated in following PABPC1 depletion are highlighted in red, while downregulated genes are highlighted in blue. Genes are labeled if they meet the log2fc and adjust p value cutoffs, and if they fall in gene ontology families related to immune genes. **E** GSEA of RNA-Seq results show pathways enriched or depleted following PABPC1 depletion. **F** DaPars analysis of RNA-seq from PABPC1 depleted B cells shows change in proximal-distal usage index (PDUI) for genes that are lengthening (red) or shortening (blue) following loss of PABPC1 (N=2). **G** Representative read density histogram of RAC1, identified by DaPars for having a higher ratio of longer 3’UTR reads following PABPC1 depletion. Reads from the PABPC1 depleted cells are show in red, while reads from vehicle controls are overlayed in blue. Blue markings below indicate annotated PAS, while red markings indicate where the poly(A) tail is added. Black bars indicate enriched PABPC1 binding windows. **H** PABCP1 enriched binding windows plotted against ΔPDUI of those genes following PABPC1 depletion. Genes that lengthen following PABPC1 depletion with PABPC1 binding sites are highlighted in red, while genes with PABPC1 binding targets that shorten following PABPC1 depletion are highlighted in blue. **I** rMATS-turbo analysis of PAPBC1 depleted Ramos B cells. Genes with more intron retention following PABPC1 depletion are displayed on the right and genes with less intron retention are displayed on the left. **J** Representative read density histogram of CD19, with the retained intron identified by rMATS-turbo highlighted.

The propensity of PABPC1 to bind the 3’UTR of its target genes (Fig 1C-D) led us to hypothesize that PABPC1 is able to globally regulate gene expression because interactions of RBPs with the 3’ end of RNA is known to change gene expression programs (52). To understand the transcriptional programs regulated by PABPC1, we used RNA-sequencing to assess differentially expressed genes following degron-mediated PAPBC1 depletion for 16 hours. Loss of PABPC1 resulted in downregulation of 798 genes and upregulation of 585 genes as quantified with DeSEQ2 (53) (Fig2C). Among the upregulated genes were the germinal center regulating cytokine IL23A (54), the FC µ receptor (FCMR), and the cytoplasmic poly(A)-polymerase, TENT5C. Due to the reported roles of IL23A and FCMR in regulating the antigen-specific immune response in B cells (55, 56), and the role of TENT5C in regulating levels of Ig during differentiation (57), we were curious about which pathways were altered following PABPC1 depletion. To that end, we performed GSEA on B cells in which PABPC1 was depleted. Noticeably, significantly enriched pathways included TNFα signaling, TgfΔ signaling, IL2/STAT5 signaling, and the inflammatory response, consistent with an increased inflammatory gene expression program (Fig 2E). Taken together, this suggests PABPC1 plays a previously unappreciated role in tuning the inflammatory response in B cells.

PABPC1 enrichment at the 3’UTR of its target genes (Fig 1C-D) also led us to hypothesize that the direct interaction led to alterations in 3’UTR length. To elucidate the regulatory role of PABPC1 on 3’ ends in B cells,we used DaPars (58) to analyze the effect of PABPC1 depletion on 3’UTR length (Fig 2F). While our data revealed limited 3’UTR changes, consistent with previous results (28), most changes were associated with longer transcripts. We examined genes that met a threshold ΔPDUI of greater than 0.2 or less than −0.2, had 30-fold coverage of the 3’UTR, and met a fold-change threshold of 0.59. Among the genes with longer relative transcripts was Rac1 (Fig 2F), a GTPase that is imperative for transducing signals through mIg that promote B cell survival (59). In neuronal contexts, the longer Rac1 isoform was associated with differentiation due to increased Rac1 signaling (60). This suggests PABPC1 may tune signaling through mIg by regulating the 3’UTR length of Rac1. Interestingly, when we correlated genes with PABPC1 binding sites to genes with changing 3’UTR lengths upon PABPC1 depletion, we found that genes to which PABPC1 bind lengthen almost exclusively when PABPC1 expression is perturbed, adding clarity to the role of PABPC1 in promoting shorter transcripts through direct interaction with the 3’UTR (Fig 2H).

In the same way APA can alter expression by lengthening the 3’ UTR, alternative splicing is another processing mechanism by which RNA transcripts can be changed to alter gene expression. PABPC1 has been implicated in alternative splicing in other systems (61), so we sought to examine the role of PABPC1 in mediating alternative splicing in B cells. Briefly, we used rMATS-turbo (62) to analyze RNA-sequencing for splicing dysregulation following PABPC1 depletion for 16hrs. Although we find few alterations in splicing, the signaling adapter CD19 showed notable intron retention upon depletion of PABPC1 (Fig 2I-J); while intron retention has not been previously attributed to PABPC1, alterations to RNA processing are likely due to indirect changes to other regulators because our CLAP data indicate PABPC1 does not bind to CD19 directly. Nevertheless, retention of this intron following PABPC1 depletion enables cancerous B cells to evade therapy (63), suggesting PABPC1 may promote survival and therapeutic resistance in B cell malignancies.

In summary, our data suggest that, globally, PABPC1 upregulates gene expression in B cells. These effects are not primarily driven by alterations to 3’UTR length or splicing. In the subset of genes that do display changes in processing, PABPC1 may help regulate signaling through mIg and drive cancer survival in a previously unrecognized way.

### PABPC1 mediated alterations in gene expression are highly context dependent

Despite numerous reports aimed at defining the function of PABPC1, the precise role it plays in cells remains ambiguous and contradictory. It has paradoxically been attributed to disparate regulatory programs leading to upregulation in some genes, and downregulation in others (27, 28, 64–67). While our global analysis revealed several alterations to gene expression and RNA processing in B cells, it was not clear if these effects were mediated by direct interactions of PABPC1 and RNA, or if these effects were indirect. Additionally, while it has been shown that PABPC1 also regulates immune gene processing in cancerous cells (68) it has not been addressed whether PABPC1 is able to directly modulate immune gene expression in B cells.

To clarify the direct effects of PABPC1 binding we asked whether the context of PABPC1 binding determines differential gene expression, drawing on both our CLAP-seq and degron-based RNA seq data. We first correlated PABPC1 binding enrichment over control with PABPC1-mediated differential gene expression on a per-gene basis (Fig 3A). Surprisingly, our analysis identified transcripts such as TNFRSF10D, FOXO3, and ALYREF with differential expression correlated with PABPC1 binding. Notably, upon PABPC1 loss, FOXO3 is upregulated, which has been shown to promote apoptosis in antigen-stimulated B cells (69), suggesting PABPC1 may restrain aberrant B cell apoptosis, further implicating PABPC1 in B cell survival following activation

**Figure 3.**
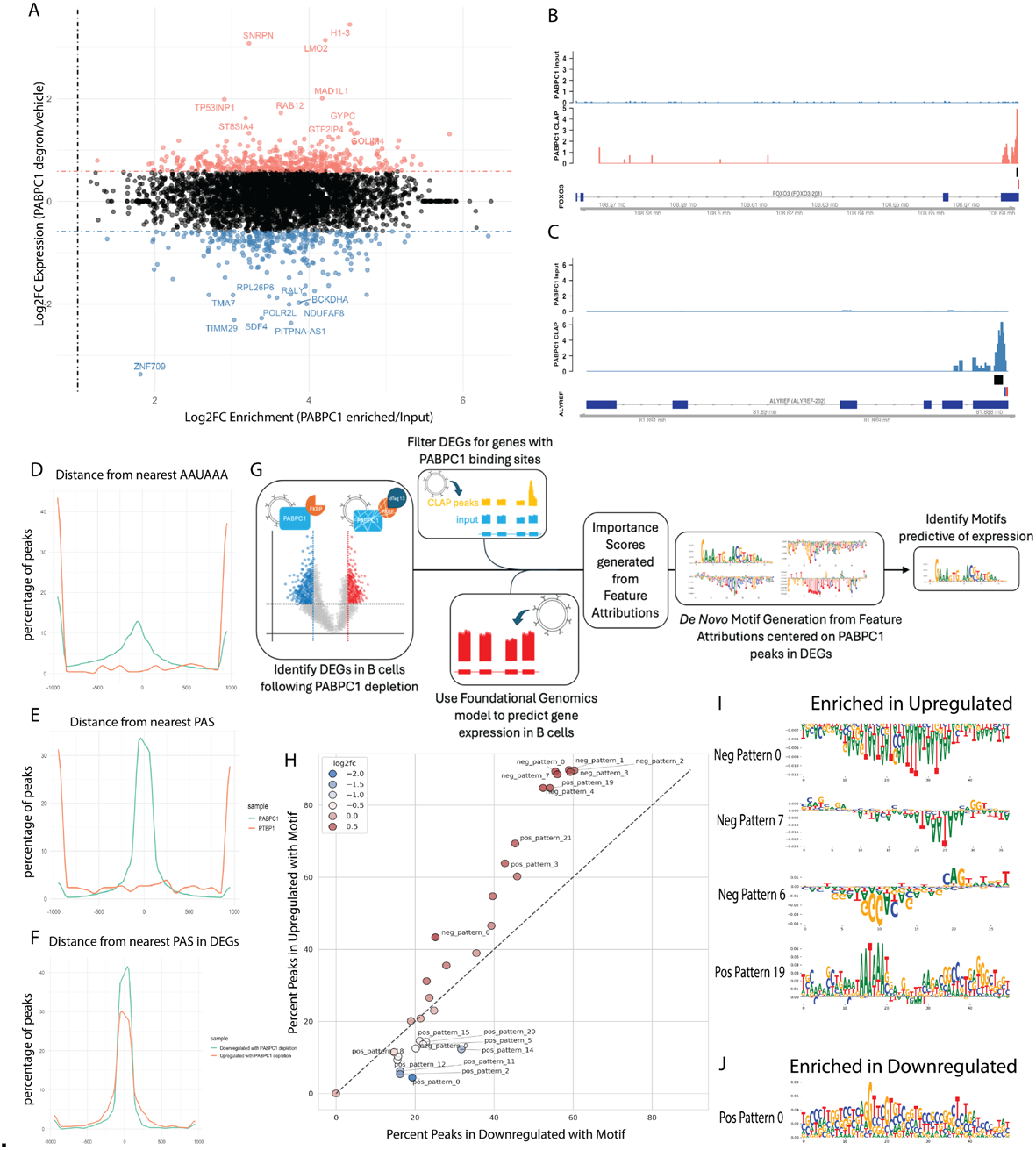
PABPC1 binding windows correlated with differentially expressed genes reveal context dependent patterns that are predictive of gene expression changes. **A** PABPC1 binding windows were correlated with genes that are up or down regulated following PABPC1 depletion. There is no correlation between binding and expression patterns. **B** Representative read density histograms of FOXO3 and (**C)** ALYREF. Both display PABPC1 enrichment in the 3’UTR, but FOXO3 is upregulated following depletion while ALYREF is downregulated. Black bars indicate PABPC1 enriched binding windows, blue bars indicate annotated PAS, and red bars indicate annotated cleavage sites **D** PABPC1 binding window distance to nearest canonical PAS or **E** any annotated PAS, with 0 centered on the PAS. PTBP1, an intronic poly-pyrimidine tract binding protein, is used as a control. **F** PABPC1 binding windows proximity to nearest PAS in upregulated and downregulated genes. **G** Schematic of Machine learning workflow. We filtered regions for differentially regulated genes following PABPC1 depletion that contained PABPC1 binding sites. A foundational genomics model was used to predict features most important for gene expression in b cells within the filtered regions. Important features are scored and motifs identified using those scores. Finally, motifs in upregulated genes are compared against binding windows within downregulated genes. **H** FIMO analysis of regions predictive of gene expression show motifs centered on a 200bp context window around PABPC1 binding sites in upregulated genes (y axis) vs downregulated genes (x axis) to determine enrichment of patterns that may explain the disparate regulatory programs mediated by PABPC1. **I** top motifs enriched in the upregulated binding windows relative to downregulated binding windows are shown. Neg Pattern is used to indicate motifs predictive of downregulation, while Pos Pattern is used to indicate motifs predictive of upregulation. **J** Regions enriched in downregulated peaks predictive of downregulation.

On a global scale, PABPC1 depletion primarily decreased gene expression (Fig 2D), indicating PABPC1 broadly promotes gene expression. However, when we focus on differentially expressed transcripts with significantly enriched PABPC1 binding sites, we notice the opposite trend; that is, 342 genes with increased expression levels and 235 with decreased expression levels. This suggests that direct interaction of PABPC1 with RNA targets is associated with repression, where loss of PABPC1 relieves inhibition of transcriptional expression.

Although there is no linear relationship between PABPC1 enrichment and magnitude of gene expression changes, this expression shift reveals a distinction between indirect and direct effects of PABPC1 binding. While It is known that PABPC1 generally balances transcript abundance by protecting the ends, it can also destabilize mRNA by recruiting cytoplasmic deadenylases that increase turnover (70, 71). Our findings support a model whereby PABPC1 exerts context dependent regulation and provide evidence that direct binding is associated with gene repression.

Our observation that PABPC1 depletion can cause both positive and negative changes in gene expression changes led us to hypothesize that context of PABPC1 binding sites underlie these paradoxical outcomes. While the primary PABPC1 motif is AAAAA in our hands, it is also able to bind directly to the PAS ‘AAUAAA’ core hexamer (37). Upon examining the motifs we identified using CLAP, we find the canonical PAS is represented by the second ranked motif from our CLAP data set (Fig1D, p1E-141), which confirms that PABPC1 interact with the PAS hexamer in B cells. To address our hypothesis, we tested whether PABPC1 differentially interacts with the PAS or other *cis*-regulatory features in a manner that could explain the discrepancy. However, it is evident that the PAS is represented in PABPC1 binding sites contained in both up and down regulated genes (Fig 3B-C), indicating that interaction with the PAS alone does not correlate with changes in gene expression.

Precise interactions in the region surrounding the PAS are required for efficient cleavage and polyadenylation, and gene expression (72). Because the PAS was present in binding sites regardless of expression changes, we reasoned that PABPC1 proximity to the PAS may influence gene expression. To address this, we sought to determine distance from the center of PABPC1 peaks to the nearest PAS. We find PABPC1 generally binds in close proximity to the PAS, as roughly 10% of peaks bound directly to the nearest AAUAAA (Fig 3D), whereas nearly 35% of peaks centered on any annotated PAS (73) (Fig 3E). As a control, we compared PABPC1 binding regions to those of the splicing regulator PTBP1, which has a well-characterized role in binding polypyrimidine tracts in introns, suggesting specificity of PABPC1 for the region surrounding the PAS. Strikingly, following PABPC1 depletion, when we compared binding regions associated with upregulated genes to those associated with downregulated genes, we find PABPC1 is more likely to be enriched within the region immediately surrounding the PAS in genes that are downregulated with loss of PABPC1 (Fig 3F). This indicates PABPC1 binding near the PAS has a positive effect on gene expression. To our knowledge this is the first report linking PABPC1-PAS proximity as a contributor to efficient gene processing.

While PABPC1 proximity to the PAS was associated with upregulation, we also took an unbiased approach at identifying additional, more subtle patterns that may help decode the PABPC1 regulatory logic. To identify cis-acting elements in the context of genes with expression programs correlated to PABPC1 binding sites, we used a foundational genomics model (Fig 3H) to identify sequences predictive of gene expression in B cells. Intriguingly, three of the top five differentially enriched motifs contained an A/U rich region that represented the canonical PAS core hexamer and was predictive of both increased and decreased gene expression (Fig 3I, Neg pattern 0, Neg Pattern 7, and Pos Pattern 19). This is supported by our observation that PABPC1 binding around the PAS was observed in both upregulated and downregulated genes (Fig3 A-C). Curiously the sole motif that was predictive of upregulation within the upregulated genes following PABPC1 loss was unique in that it alone contained a GC-rich region immediately downstream of the PAS. This suggests that interactions with the context window surrounding the PAS is the primary predictor of gene expression.

Our observation that a downstream G-rich element is predictive of gene expression is consistent with known 3’ end processing mechanisms. CSTF-64 is an indispensable core member of the CPA machinery that is known to bind in the G-rich region immediately downstream of the PAS to recruit other CPA factors to the PAS site (74– 76), and is implicated to regulate APA in B cells (10, 11). To lend support to the accuracy of our model predictions, Pos Pattern 19 was remarkably similar to the second most significantly enriched motif identified by Homer in our CLAP-seq data (Fig 1E). This is consistent with a model in which PABPC1 may suppress genes containing this motif by directly binding to the downstream G-rich element to sterically block efficient 3’ end processing. Strikingly, the most enriched motif within PABPC1 binding windows in the downregulated genes following PABPC1 depletion relative to the upregulated genes was a degenerate G/C rich sequence. While only enriched in 20% of the downregulated genes, it suggests PABPC1 may be able to interact with G/C-rich regions outside of its canonical binding preferences, a finding that corroborates early reports that the PABPC1 is able to bind non-poly(A) regions (41, 77). Taken together, our results highlight the binding context of PABPC1 as a key regulatory determinant of gene expression.

### PABPC1 preferentially binds the sIg 3’UTR to regulate IgM expression

Whether Ig is expressed on the surface of a cell or secreted is dictated primarily by APA (7–9). We sought to determine the role PABPC1 plays in regulating Ig expression because it is a known regulator of APA (28, 29, 78) and has been previously correlated with Ig secretion (15). Analysis of the binding sites for PABPC1 in our CLAP-seq dataset revealed PABPC1 binds to the 3’UTR of both sIg and mIg (Fig 4A). Using peak area in region 100bp upstream of each respective PAS, we reveal PABPC1 is more significantly enriched at the sIg PAS than the mIg PAS (p=0.0376), indicating PABPC1 binds preferentially to the sIg isoform (Fig 4B). Where others have correlated PABPC1 binding to IgM, this study supplies the first evidence that PABPC1 directly interacts with Ig RNA at the sIg PAS

**Figure 4.**
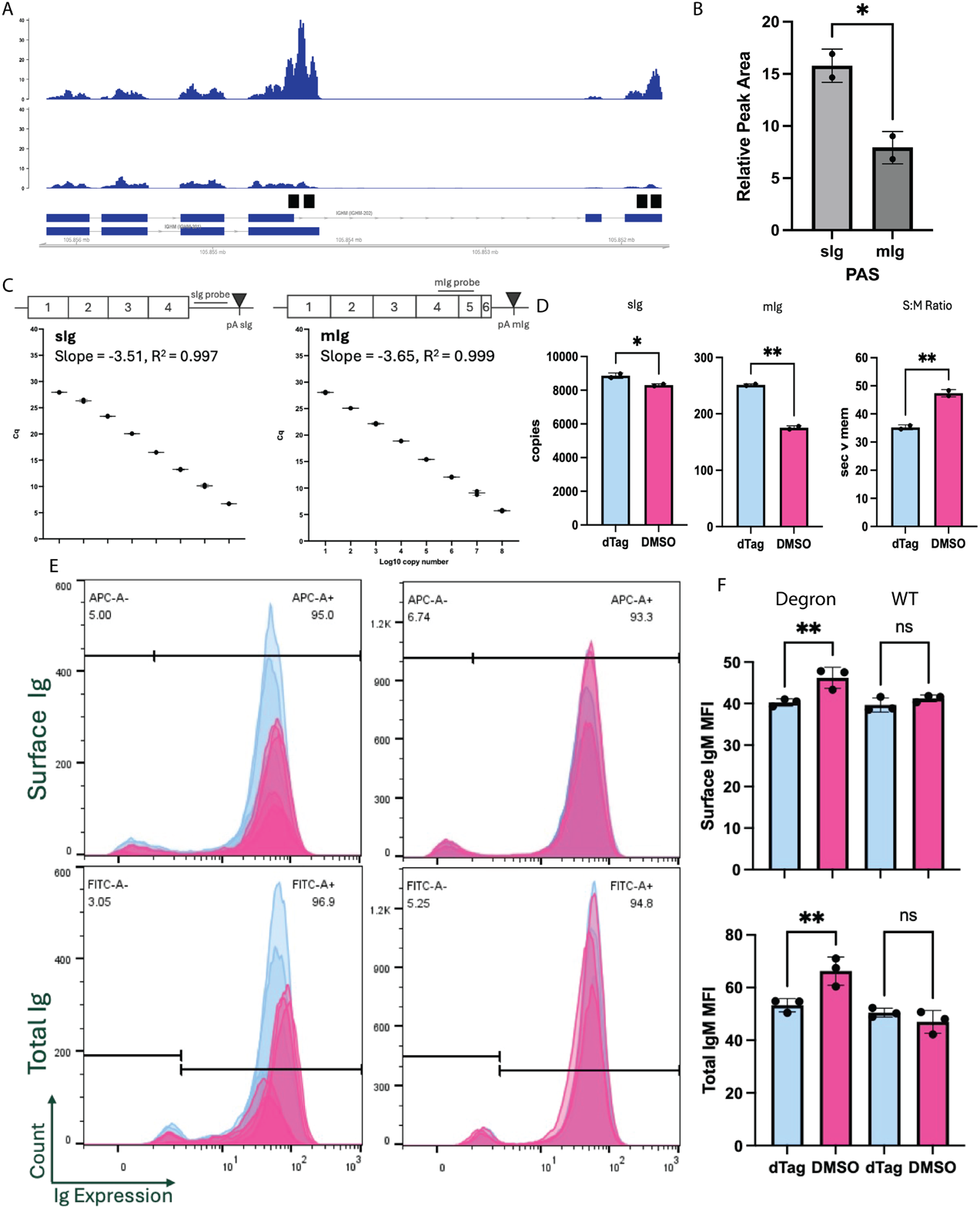
PABPC1 modulates IgM expression through direct interaction with the 3’UTR. **A** Read density histograms of CLAP reads (top) and input reads (bottom) for PABPC1 enrichment over the Ighm gene. Black boxes denote regions that were identified for significantly enriched binding over background. **B** Area under the curve was calculated from read density histograms normalized using transcripts per million for the region 100bp upstream of the PAS and input subtracted from CLAP samples. PABPC1 is more significantly enriched at the sIg PAS region (n=2, p=0.0376). **C** Standard curves for determining absolute copy number of each isoform using qPCR. Probe locations are schematized above each respective plot. **D** RT-qPCR for absolute copies of sIg (left, n = 2, p = 0.0457) mIg (middle, n = 2, p=0.0013) and change in sIg to mIg ratio (right, p = 0.0084) in dTag13 treated (and thus PABPC1 depleted) vs DMSO vehicle controls. **E** Intracellular Flow cytometry shows representative histograms of IgM on the surface (top, p = p = 0.0043) or total IgM expressed by cells (bottom, p = 0.006) in dTag13 treated cells vs DMSO control (N=3). **F** Quantification of MFI for plots shown in (E).

While we show evidence that PABPC1 binds directly to IgM transcripts, function cannot be inferred from binding alone. To determine if PABPC1 expression had any functional consequences on Ig expression, we utilized our degron system. We designed qPCR probes to measure the absolute copies of both sIg and mIg transcripts (Fig 4C). Depletion of PABPC1 modestly increased copies of the sIg 3’UTR by 552.6 (p = 0.0457) while it increased copies of mIg by 76 (p = 0.0013). Thus, depletion decreased the ratio of sIg:mIg by 25% due to the greater fold increase of mIg relative to sIg, suggesting PABPC1 promotes the usage of the proximal PAS (Fig 4D), in line with the binding preferences displayed in our CLAP dataset (Fig 1C, 4A) and the general trend in 3’UTR changes displayed by DaPars analysis. At the post-transcriptional level, our data confirm the reported role PABPC1 plays in determining antibody secretory status (15), and further offers direct evidence that PABPC1 interacts with Ig RNA.

Protein expression typically correlates with RNA expression. Therefore, considering the increase in abundance of both sIg and mIg transcripts, we expected to observe a corresponding increase of Ig protein levels. To determine the change in sIg:mIg ratio following depletion, B cells were stained for mIg, fixed and permeabilized, and stained for total Ig following PABPC1 depletion (Fig 4E). Changes in mean fluorescence intensity (MFI) can then be used as a metric for changes in amount of protein. Surprisingly, we observed that PABPC1 depletion caused a decrease in mIg MFI by nearly 6 (p = 0.0043) compared to 1.6 in the DMSO vehicle control (p = 0.459), while depletion also caused a decrease in total Ig MFI by 12.97 (p=0.006), compared to a small but insignificant (p=0.4955) increase in MFI by 3.5 in the DMSO vehicle control (Fig 4F). Taken together this suggests PABPC1 promotes translation of Ig transcripts in a manner decoupled from its regulation at the post-transcriptional level.

## Discussion

Our findings are consistent with other studies undertaking depletion of PABPC1 using knockdown in a variety of cell types, where PABPC1 depletion resulted in altered gene expression. In polyadenylated transcripts, PABPC1 binding is associated with less turnover because it is able to protect the 3’ end from degradation (79). However, paradoxically, it also recruits cytoplasmic deadenylases that work to degrade the poly(A) tail (70). Mechanistically, these competing paradigms have not yet been reconciled by our molecular understanding of PABPC1 binding, but it stands to reason that multiple sequences might encode different functions of PABPC1 or gene body context might encode these differences. Nevertheless, we show PABPC1 depletion leads to differential gene expression in inflammatory immune genes, though displays few changes in 3’UTR length. This is consistent with the observations of Kowalski et al., which established that knockdown of PABPC1 led to changes in RNA abundance, but had few consequences on 3’UTR length. While we see a small effect of PABPC1 perturbation on 3’UTR length, our results are directly contradictory to Li et al, who reported PABPC1 promotes lengthening transcripts. The few genes that displayed altered 3’UTR length suggest PABPC1 actually promotes shortening of transcripts. This may be due to regulatory differences between muscle cells and immune cells and highlights the context specific effects on 3’ end formation mediated by PABPC1.

The enigmatic consequence of PABPC1 perturbation has confounded the literature, and in our study, binding yields both up and down regulation. One way to conceptualize the multifaceted roles of PABPC1 is to suggest that binding context determines function. In the way that HNRNP and SR proteins binding location determines outcomes in a context specific way (80, 81), here too, we found motifs correlating with up and down regulation of gene expression using deep learning. One mechanistic way this might occur is that other proteins nucleate with PABPC1 to confer these heterogeneous outcomes. Another possibility is that PABPC1 may compete with other RBPs for similar binding motifs. We note that the third ranked motif identified in enriched PABPC1 binding windows is identical to the binding site for the Pumilio family of RBPs (82–84) (Fig 1D), suggesting that PABPC1 may exert indirect influences by competing with PUM1 or PUM2. In support of this, PABPC1 overexpression is reported to inhibit PUM-mediated repression of RNA targets (51).

Previous reports that perturbed PABPC1 in B cells found a decrease in mIg but no change in total Ig, suggesting PABPC1 was influencing Ig expression(15). Our data indicate PABPC1 depletion does indeed decrease mIg, though they also show a decrease in total Ig, in line with the well documented role of PABPC1 in facilitating translation. It’s possible that these disparities are driven by how PABPC1 depletion is achieved. While the degron system we favored allows acute control of PABPC1 expression, others have chosen constitutive expression of PABPC1 siRNA (15, 28, 78) which may cause the cells to develop compensatory mechanisms following long-term loss of PABPC1. Additionally, while Peng et al establish a correlative link between PABPC1 and the sIg 3’UTR, it could not be ruled out if the interaction was mediated by the poly(A) tail, considering the canonical binding motif for PABPC1 is poly(A). Here, we establish that the interaction is sequence specific by biochemically mapping PABPC1 directly to both the sIg and mIg 3’UTR. While our data show a non-significant trend towards the sIg 3’UTR based on normalized read counts, Peng et al show a preference for the sIg 3’UTR over mIg using RIP-qPCR. Importantly, perturbation does not fully switch B cells to a secretory phenotype, indicating that there may be other factors that directly mediate the switch from mIg to sIg. Nevertheless, taken together, our data lend further evidence to support the role for PABPC1 in modulating the delicate interplay between mIg and sIg to help shape humoral immunity.

To map where RBPs bind, several similar sequencing-based methods have been developed that take advantage of the ability of UV light to induce photo-crosslinks between proteins and RNA in direct contact (33, 34, 85–90). Indeed, others have employed CLIP-seq to determine where PABPC1 binds in MEL cells (32). While CLIP-seq allows capture of endogenous RNA-protein interactions, the immunoprecipitation-based strategy has several downsides. Affinity of antibody for its target is heterogenous between specific targets and, despite possessing generally high affinity, the interaction is sensitive to highly stringent wash conditions, which means more gentle washes could potentially lead to increased noise. We have chosen to use CLAP-seq, which employs an overexpression system with HaloTagged constructs to take advantage of the covalent interaction between Halo and its ligand. This helps to decrease noise at the cost of losing endogenous interactions, but we have sought to mitigate overexpression artifacts by using a doxycycline-inducible strategy. Importantly, when comparing our data to previously described CLIP-seq experiments that determined PABPC1 binding sites in MEL cells, we find a high level of concordance between the two datasets, both in terms of propensity to bind 3’UTR, and sequence specificity (32) (Fig 1B, D). While our data were obtained from human cells, the conserved binding patterns of PABPC1 in the two cell lines and the sequence conservation of PABPC1 across species highlights its cellular importance. Importantly, PAPER-CLIP is a method to isolate poly-Adenylated RNA on the basis of PABPC1s affinity for the poly(A) tail. However, given the ability of PABPC1 to bind to other regions of RNA besides the poly(A) tail, both reported by kini et al and displayed by our data, this suggests careful consideration must be used when determining poly(A) enrichment strategies.

PABPC1 regulation might just be beginning to be understood. The essential role for PABPC1 in the cell makes perturbation studies to determine function difficult and remains a limitation of this study, despite adopting the degron system to mitigate the negative effects of constitutive perturbation. Additionally, other cytoplasmic PABs have been shown to compensate for loss of PABPC1 in other systems (27, 49). It’s possible that upregulation of highly homologous PABPs limit the mechanistic insights that can be gained from perturbation. Additionally, PABPC1 has been reported to have pleiotropic functions in the cell, including reported roles in translation, RNA trafficking, RNA stability, and paradoxically, RNA de-adenylation and turnover. The many cellular roles ascribed to PABPC1 has made it difficult to decouple its effects on gene expression and translation. Here, we begin to provide clarity into the post-transcriptional regulatory mechanisms controlled by PABPC1 and suggest that they are highly context dependent. Understanding the rules of that context dependence will be the key to decoding the role of PABPC1 in sculpting the post-transcriptional landscape of B cells.

## Data Availability

The computational pipeline used to identify sequence features and calculate per-nucleotide attributions associated with PABPC1-dependent mRNA regulation is openly available. The core analysis scripts, including the configurations for Borzoi model execution and motif enrichment, are archived on Zenodo (DOI: 10.5281/zenodo.19154687). To ensure full reproducibility of the pre-computed attribution handling and de novo motif discovery described in this manuscript, the specific modified software dependencies utilized in this study have also been archived:

- A modified fork of the genomics deep learning library gReLU (95), containing the functional updates for pre-computed motif attribution mapping, is available on Zenodo (DOI: 10.5281/zenodo.19154702).
- The corresponding modified version of tfmodisco-lite, utilized for sequence motif discovery, is available on Zenodo (DOI: 10.5281/zenodo.19154712).

All custom code, configuration files, and environments required to reproduce the foundational genomics analyses and generate the figures presented in this study are included in the primary repository.

The custom scripts used to generate all other data and figures presented in this study are also stored on Zenodo (DOI: 10.5281/zenodo.19188093).

Sequencing tracks for CLAP-seq experiments can be viewed with the UCSC genome browser session URL: https://genome.ucsc.edu/s/zdmiller21/PABPC1_CLAP. Corresponding data is stored on the Gene Expression Omnibus with the accession number GSE326352 (reviewer token: kjqhokosbpgtfmj) Sequencing tracks for RNA-seq experiments using degron-mediated depletion of PABPC1 can be viewed with the UCSC genome browser session URL: https://genome.ucsc.edu/s/zdmiller21/PABPC1_degron. Corresponding data is stored on the Gene Expression Omnibus with the accession number GSE326353 (reviewer token: sjsxoqumxjydveb).

## Supplementary Data Statement

Supplementary Data are available at NAR Online.

## Acknowledgements

We thank Jon Boyson for his valuable guidance and support. We acknowledge the use of both the Oklahoma Medical Research Foundation sequencing core, and the University of Vermont Integrative Genomics Resource Core Facility (RRID:SCR_021775). We also acknowledge the use of AI in generating custom scripts and analyzing sentence structure to identify text to modify during manuscript revision. The schematic depicted in Figure 1A was created in BioRender. Miller, Z. (2026) https://BioRender.com/zsc7f80.

---

**Supplementary Table 1.**
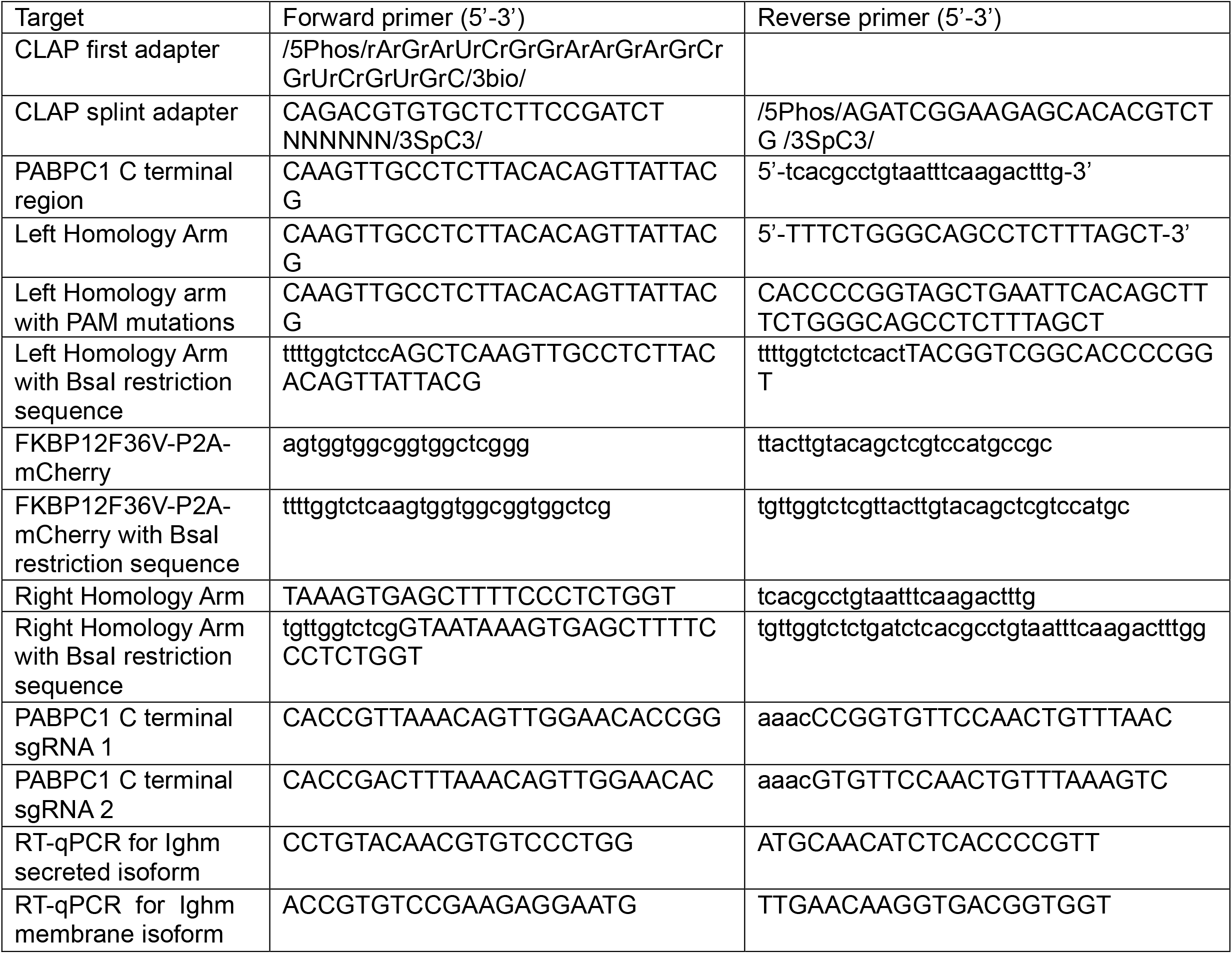
All primers used for library preparation, generation of the PABPC1 homology directed repair template, sgRNA sequences, and RT-qPCR during the course of this study.

